# Hierarchical temporal transformer for cancer grade prediction and cross-cancer transfer learning from pathology reports

**DOI:** 10.64898/2026.09.09.750492

**Authors:** Nura Brimo, Rajneesh Anand, Hassan Harb, Dilek Cokeliler Serdaroglu

## Abstract

Language models for cancer clinical reports carry two blind spots. They read each report in isolation, ignoring how a patient’s disease changes across visits, and they are evaluated only on cancer types present in their training data. We present the Hierarchical Temporal Transformer (HTT), a two-level architecture that addresses both. Level 1 encodes each report with BiomedBERT adapted by low-rank adaptation (LoRA). Level 2 is a temporal transformer that reads a patient’s full report sequence using a continuous-time positional encoding built from the measured number of days between visits, with learnable cancer-type embeddings supplying per-family conditioning. Two experiments test the two capabilities separately, since no fully open corpus contains longitudinal reports for many cancer types. On a controlled synthetic corpus of sequential radiology reports, in which progression phrases are inserted from templated trajectories, HTT reaches a validation AUROC of 0.942 against 0.881 for a single-report baseline and transfers to held-out pancreatic cancer at 0.995 against 0.949, while the two models are indistinguishable on a 60-patient test set. On 4,786 real pathology reports from the TCGA-Reports corpus spanning 14 cancer types, HTT predicts tumor grade for three types withheld entirely from training, reaching AUROC 1.000 on thyroid carcinoma, 0.960 on sarcoma and 0.808 on lung squamous cell carcinoma. The mean held-out AUROC of 0.923 equals the in-distribution test AUROC of 0.923, so transfer to unseen cancer families incurred no measurable penalty. Ablation on the real corpus shows that the transfer is carried by the pre-trained encoder rather than by the temporal components, which, with one report per patient, contribute 0.39 AUROC points. Grade-related pathological language therefore appears to be learnable in a cancer-type-agnostic way, which points toward unified cancer NLP systems that require no per-type retraining.

## 1 Introduction

Cancer care produces text at every stage. Radiologists write imaging impressions, pathologists describe resected tissue, and oncologists synthesize both in clinical notes. Models that read these documents have proved useful for predicting progression from radiology impressions [1,2], extracting phenotypes and generating diagnoses [3], and classifying disease response [4]. Like clinical language models more generally [5], they share two limitations that this study is designed to address.

The first is the absence of temporal modelling. Every prior system we reviewed treats each clinical report as an isolated document: the model reads one report and predicts from that report alone. Patients, however, are seen repeatedly over months or years, and the shape of that history carries information. A patient whose scans move from stable disease, to enlargement, to new nodal involvement is in a different situation from one whose most recent scan alone mentions new spread, even if the final reports read identically. A model that sees only the last report cannot distinguish them.

The second is the absence of cross-cancer-type transfer. A comprehensive cancer center treats dozens of tumor types, and building and maintaining a separate model for each scales poorly. For rare cancers the problem is not cost but feasibility: an institution may never accumulate enough labeled records to train on at all [6]. Prior work either trains and tests on a single type or, where several types are used, evaluates only on types seen during training. No prior study asks whether a model trained on breast, lung and kidney cancer can read a thyroid or sarcoma report it has never encountered.

We developed the Hierarchical Temporal Transformer (HTT) to address both. Level 1 uses BiomedBERT [7], a biomedical language model pre-trained on PubMed abstracts and PubMed Central full text, adapted with LoRA [8] so that 11.1% of parameters are trainable, to encode each report as a dense vector. Level 2 is a temporal transformer [9] that consumes the sequence of a patient’s report vectors together with the actual number of days elapsed between visits, and conditions its reasoning on a learnable cancer-type embedding so that one set of weights can serve many tumor families.

The two capabilities are tested in separate experiments, because no fully open corpus contains longitudinal clinical reports for many cancer types without a data-use agreement. Experiment 1 evaluates temporal modelling on a controlled synthetic corpus of sequential radiology reports, which permits a clean comparison against a single-report baseline. Experiment 2 evaluates cross-cancer transfer on the TCGA-Reports corpus [10,11], comprising 4,786 grade-labeled real pathology reports across 14 cancer types, with three types withheld entirely from training.

We ask three questions. Does reading the full visit sequence with real elapsed time improve prediction over reading the most recent report? Does tumor grade prediction transfer to cancer types absent from training? And when transfer is observed, which component carries it? Grade is the right target for the second question because it is defined for essentially all solid tumors, drives treatment intensity, and is expressed in language that differs between cancer families. Thyroid carcinoma is graded through mitotic activity, tumor necrosis, aggressive variant histology and invasion-related findings such as extrathyroidal extension. Sarcoma uses the FNCLCC system, a composite of mitotic count per high-power field, percentage necrosis and a differentiation score, whose format appears nowhere in a carcinoma corpus. The biological concept is shared; the surface language is not.

## 2 Contributions and comparison with prior work

This work provides, to our knowledge, the first demonstration that pathological grade language transfers across biologically distinct cancer families under strict held-out conditions. A mean AUROC of 0.923 on three cancer types entirely absent from training, equal to in-distribution performance, indicates that the model has learned a representation of pathological aggressiveness rather than type-specific surface features.

Three elements are new. First, a continuous-time positional encoding for clinical report sequences. Prior temporal models in NLP assign integer positions 1, 2, 3 to sequential items, which implicitly assumes equal spacing. Clinical visits are irregular; a scan two weeks after surgery means something different from one six months later. We apply sinusoidal encoding to the actual elapsed days, with learnable frequency scaling that lets the model discover clinically relevant time scales during training. Second, a systematic held-out-type evaluation of free-text cancer report understanding, on 459 real reports from three cancer types never seen in training. Third, a hierarchical design that separates report-level encoding, which is parameter-efficient, from patient-level temporal reasoning, which retains full capacity. Table 1 places the work against prior systems.

**Table 1.** Comparison with prior cancer NLP studies. AUROC values for prior work are approximate, read from published figures.

| Study | Data | Types | Temporal | Unseen-type eval. | Best AUROC |
| --- | --- | --- | --- | --- | --- |
| Woollie [2] | Radiology reports | 1 | No | No | 0.80 |
| CancerLLM [3] | Clinical notes | Multiple | No | No | 0.85 |
| Tan et al. [4] | Radiology reports | 1 | No | No | 0.82 |
| Lee et al. [12] | Image and text | Multiple | No | Partial | 0.88 |
| HTT (this work) | Pathology reports | 14 (3 held out) | Yes | Yes | 0.923 (mean) |

## 3 Methods

### 3.1 Study design

Figure 1 gives the overall design. Two experiments share one model and one training procedure but use different corpora and different labels. Experiment 1 uses synthetic sequential radiology reports to test whether reading a patient’s full visit history improves progression prediction over reading the most recent report alone. Experiment 2 uses real TCGA pathology reports to test whether grade prediction transfers to cancer types withheld from training. In both, a non-temporal Baseline sharing the same encoder and LoRA configuration is trained under identical conditions, so that any difference isolates the contribution of the temporal and conditioning components rather than of the pre-trained encoder.

**Figure 1.**
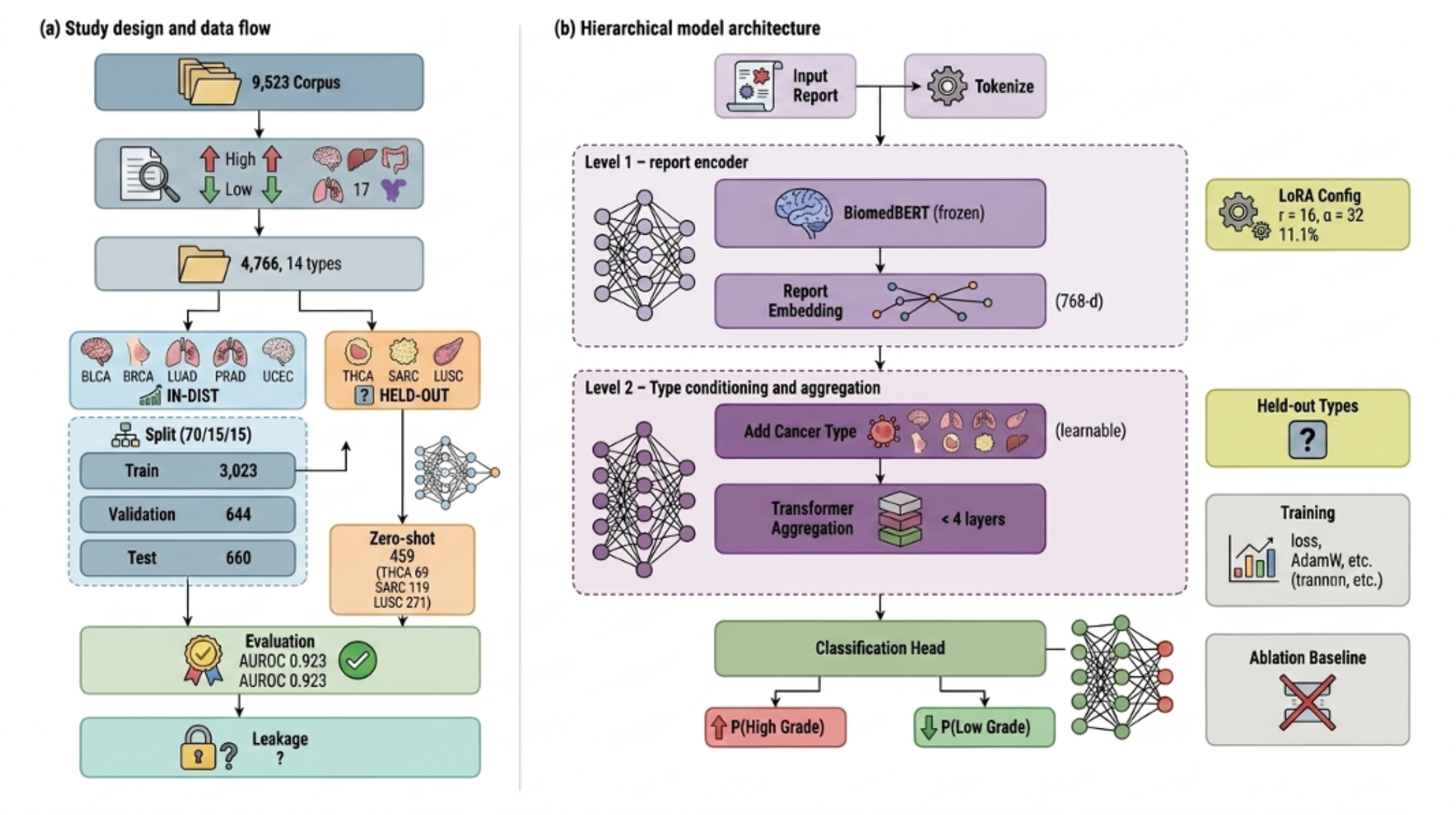
Study design and model architecture. (a) Experiment 1 uses a synthetic corpus of sequential radiology reports across four training cancer types with pancreatic cancer held out; the label is progression at any visit. Experiment 2 uses the TCGA-Reports corpus with rule-based grade labels across 11 training types and three withheld types; the label is high against low grade. In both, the full model and a non-temporal Baseline are trained identically and scored by a single checkpoint. (b) Each report, prefixed with a structured cancer-type token, is encoded independently by BiomedBERT with LoRA, mean-pooled and projected to 512 dimensions (Level 1). The sequence of report embeddings receives a continuous-time positional encoding computed from elapsed days and a learnable cancer-type embedding, then passes with a prepended CLS token through a four-layer Transformer encoder (Level 2). A patient-level head on the CLS output carries the primary loss; a visit-level head on each visit output supplies an auxiliary loss during training only. The Baseline reads the most recent report alone and classifies from the Level 1 embedding through a single linear layer.

### 3.2 Data

#### Synthetic sequential radiology reports (Experiment 1)

No publicly available corpus contains multiple longitudinal radiology or pathology reports per cancer patient without a data-use agreement, so a syn-thetic corpus was generated programmatically. The generator produces free-text radiology impressions with realistic anatomical language, measurement patterns and disease trajectories. Cohorts comprised 500 patients across five cancer types: 100 each of non-small cell lung cancer (NSCLC), breast cancer, colorectal cancer (CRC) and prostate cancer for training, plus 100 pancreatic cancer patients withheld entirely for held-out evaluation.

Each patient was assigned one of four longitudinal trajectories: stable at every visit (30%); progression beginning at the midpoint of the sequence (30%); progression beginning only in the final two visits (25%); and initial response followed by late relapse (15%). Patients received between 2 and 8 visits, with intervisit intervals drawn uniformly from 42 to 90 days, giving the irregular spacing characteristic of real scheduling. Start dates were randomized across 2018 to 2022, and the days elapsed from the first visit were recorded per visit and fed to the continuous-time encoding.

At each visit, one of four type-specific radiology templates was selected at random, matched to the modality used for that cancer: CT chest for NSCLC, contrast-enhanced breast MRI, CT abdomen and pelvis for CRC and pancreatic cancer, and multiparametric prostate MRI. Variable slots (lesion size, comparison size, nodal status, and modality-specific values such as SUVmax or PI-RADS score) were filled with plausible sampled values. The disease-change slot was filled deterministically from the assigned trajectory: progressing visits received phrases such as “enlarged”, “interval increase” or “progressive dis ease”, and non-progressing visits “unchanged”, “stable in size” or “no significant interval change”. A structured cancer-type token was prepended to every report. All random operations were seeded at 42.

Because the progression label is written into the text by construction, this corpus does not test whether the model can infer progression from subtle clinical language. It tests something narrower and still informative: whether an architecture that reads the whole sequence with real elapsed time can integrate trajectory information that a single-report model cannot access, in a setting where the ground truth is known exactly. Visit-level labels (1 = progressing at this visit) were assigned from trajectory and visit index. The patient-level label is the maximum across visits, so a patient is a progression case if progression occurred at any visit.

#### TCGA-Reports (Experiment 2)

TCGA-Reports [11] is a fully open corpus of 9,523 free-text pathology reports from The Cancer Genome Atlas [10], hosted on the Hugging Face Hub with no data-use agreement. Each record pairs a patient identifier with a report written by a pathologist after surgical resection or biopsy, covering histology, grade, differentiation, tumor size, margin status and nodal involvement. The corpus contains one report per patient, so in Experiment 2 the temporal component operates on sequences of length one. The two capabilities of the model are therefore necessarily evaluated on separate corpora, a constraint we return to in the Limitations.

The dataset provides no structured cancer-type labels. We extracted cancer site directly from report text using 17 ordered pattern sets over anatomical and histological keywords, applied most-specific-first so that squamous lung reports are not absorbed by the general lung pattern. Table 5 in the appendix Table 2 gives representative patterns. Retaining types with at least 40 labeled reports left 14 cancer types.

**Table 2.** Representative cancer-site extraction patterns for selected TCGA cancer types, applied case-insensitively and most specific first.

| Code | Cancer type | Representative patterns |
| --- | --- | --- |
| LUSC | Lung squamous cell | squamous.*lung, lung.*squamous |
| LUAD | Lung adenocarcinoma | \blung\b, pulmonary |
| BRCA | Breast cancer | \bbreast\b |
| KIRC | Kidney renal clear cell | \bkidney\b, renal cell |
| SARC | Sarcoma | leiomyosarcoma, \bsarcoma\b |
| THCA | Thyroid carcinoma | \bthyroid\b |
| GBM | Glioblastoma | glioblastoma, \bglioma\b |

Splits were constructed at the patient level. The 11 in-distribution types were divided 70/15/15 into training (3,023 reports), validation (644) and test (660) with a fixed seed. The three withheld types form a single evaluation set of 459: 69 thyroid carcinoma, 119 sarcoma and 271 lung squamous cell carcinoma. Held-out evaluation draws no training or model-selection samples from those types by definition, and subsampling them would only widen confidence intervals that are already wide. Table 3 lists all splits.

**Table 3.** Dataset split sizes for both experiments.

| Experiment | Unit | Train | Validation | Test | Held out |
| --- | --- | --- | --- | --- | --- |
| 1 (synthetic) | Patients | 280 | 60 | 60 | 100 (pancreatic) |
| 2 (TCGA) | Reports | 3,023 | 644 | 660 | 459 (THCA 69, SARC 119, LUSC 271) |

### 3.3 Preprocessing and labeling

#### Grade labels for TCGA

The task is binary: high-grade (label 1) against low-grade (label 0). High-grade patterns target “grade III”, “grade 3”, “poorly differentiated”, “undifferentiated”, “high grade”, “nuclear grade III”, “histologic grade III” and “FNCLCC grade 3”. Low-grade patterns target “grade I”, “grade 1”, “well differentiated”, “low grade”, “histologic grade I” and “FNCLCC grade 1”. A report matching only high-grade patterns is labeled 1 and one matching only low-grade patterns is labeled 0. Reports matching both, usually mixed-grade tumors or comparisons across several specimens, are labeled 1, since the overall grade is conventionally taken from the highest-grade component. Reports matching neither were excluded. Of 9,523 reports, 4,737 carried no grade keywords and were dropped, leaving 4,786 labeled reports. The class ratio is roughly 2.5:1 high to low (3,428 against 1,358), a mild imbalance addressed by taking AUROC, which is insensitive to class ratio, as the primary metric.

#### Cancer-type tokens

A structured token is prepended to each report before tokenization. For TCGA the token uses the project code: the 11 training types are BLCA (bladder urothelial carcinoma), BRCA (breast invasive carcinoma), COAD (colon adenocarcinoma), GBM (glioblastoma multiforme), HNSC (head and neck squamous cell carcinoma), KIRC (kidney renal clear cell carcinoma), LIHC (liver hepatocellular carcinoma), LUAD (lung adenocarcinoma), PRAD (prostate adenocarcinoma), STAD (stomach adenocarcinoma) and UCEC (uterine corpus endometrial carcinoma); the withheld types are THCA, SARC and LUSC. For the synthetic corpus the tokens are [CANCER:LUNG], [CANCER:BREAST], [CANCER:COLORECTAL], [CANCER:PROSTATE] and [CANCER:PANCREATIC]. The token supplies an explicit textual cue and maps to an integer index consumed by the type-embedding layer. Indices are assigned by alphabetical enumeration and carry no ordinal meaning.

### 3.4 Model architecture

HTT comprises a Level 1 report encoder, a Level 2 temporal transformer and two classification heads. It is implemented in PyTorch with the Hugging Face Transformers and PEFT libraries.

#### Level 1: report encoder

Each report is encoded independently by BiomedBERT-base-uncased, a 110M-parameter BERT-family model pre-trained from scratch on PubMed abstracts and PubMed Central full text [7]. It was chosen for its biomedical pre-training and open availability; BioGPT-Large (1.5B parameters) exceeded available VRAM at usable batch sizes. LoRA [8] adapts the query and value projections of every attention layer with rank *r* = 16, scaling *α* = 32 and dropout 0.1, leaving roughly 11.1% of parameters trainable. Reports are truncated to 256 tokens, which covers more than 95% of both corpora. The last hidden state is mean-pooled over non-padding tokens into a 768-dimensional report embedding and projected to 512 dimensions through a Linear, LayerNorm and GELU block. At inference, all reports in a patient’s sequence are encoded in parallel by flattening the batch and sequence dimensions.

#### Level 2: temporal transformer

Three operations are applied in sequence to the 512-dimensional report embeddings.

##### Continuous-time positional encoding

Elapsed time since the first visit, normalized to years, is encoded by sinusoidal functions over a bank of 256 frequency channels, producing a 512-dimensional time vector per visit that is added elementwise to the report embedding. Unlike standard positional encodings built on integer indices, this uses measured calendar time. Each frequency is scaled by a learnable parameter, so the model can discover which time scales, from days to years, are most predictive.

##### Cancer-type conditioning

A learnable 512-dimensional embedding is defined for each cancer type in the extraction vocabulary, plus one reserved index for types absent from training. The embedding for the patient’s type is added to every visit embedding, conditioning the whole sequence so that one set of transformer weights can behave differently for a breast, kidney or sarcoma patient. For held-out types, the prepended text token provides type-specific signal at Level 1, partially compensating for the absence of a trained embedding.

##### Transformer encoder

A learnable CLS token is prepended and the sequence passes through four Transformer encoder layers [9] with 8 attention heads, feed-forward dimension 2,048 and pre-LayerNorm. A padding mask prevents attention to padded visit positions. The CLS output is the patient-level representation; the remaining outputs, one per real visit, are the visit-level representations.

#### Classification heads

The patient-level head takes the CLS vector through Dropout(0.2), Linear (512*→*256), GELU, Dropout(0.2) and Linear (256*→*2), producing two logits from which probabilities are obtained by SoftMax. The visit-level head applies Dropout(0.2) and Linear (512*→*2) to each visit output and is used only during training to compute an auxiliary loss.

#### Baseline

The Baseline shares Level 1 exactly, down to the same BiomedBERT weights, LoRA configuration and projection layer. It differs in one respect: it processes only the most recent report for each patient, discarding all prior visits, and passes that single embedding through Dropout(0.2) and Linear (512*→*2). It has no temporal transformer, no time encoding and no cancer-type embedding. This mirrors the single-report design of prior systems such as Woollie [2] and CancerLLM [3]. Hyperparameters are identical to HTT, so any performance gap reflects the temporal and conditioning components alone. Table 4 lists the two side by side.

**Table 4.** Architectural comparison between HTT and the Baseline.

| Component | HTT (full model) | Baseline (ablation) |
| --- | --- | --- |
| Report encoder | BiomedBERT + LoRA (shared) | BiomedBERT + LoRA (shared) |
| Reports used per patient | All visits in sequence | Most recent visit only |
| Time encoding | Continuous time (days) | None |
| Cancer-type conditioning | Learnable embedding | None |
| Temporal transformer | 4 layers, 8 heads | None |
| CLS aggregation | Yes | No |
| Classification head | 2-layer MLP | Single linear layer |
| Trainable parameters | 11.1% of total | 11.1% of total |

**Table 5.** Full hyperparameter list, shared across both experiments.

| Hyperparameter | Value | Rationale |
| --- | --- | --- |
| Encoder | BiomedBERT-base-uncased | Biomedical pre-training; fits 8 GB VRAM |
| Max report length | 256 tokens | Covers >95% of reports |
| Max sequence length | 10 visits (synthetic), 1 (TCGA) | Matches corpus structure |
| Hidden dimension | 512 | Capacity against memory |
| Temporal layers | 4 | Standard for moderate sequences |
| Attention heads | 8 | Divisor of 512 |
| Feed-forward dim | 2,048 | $4 \times$ hidden dimension |
| LoRA rank ($r$) | 16 | Standard setting |
| LoRA alpha | 32 | $2 \times$ rank |
| Learning rate | $2 \times 10^{-4}$ | Selected on validation |
| Batch size | 8 (effective 32) | GPU memory constraint |
| Gradient accumulation | 4 steps | Simulates larger batch |
| Auxiliary loss weight | 0.3 | Encourages visit-level learning |
| Dropout | 0.1 (temporal), 0.2 (classifier) | Regularization |
| Max epochs | 20 | Sufficient for convergence |
| Early stopping | Patience 5 | Prevents overfitting |
| Optimizer | AdamW | Decoupled weight decay |
| Weight decay | 0.01 | Standard L2 strength |
| LR warmup | 10% of steps | Stabilizes early training |
| Mixed precision | fp16 AMP | Memory and speed |
| Random seed | 42 | Reproducibility |

### 3.5 Training

Two cross-entropy losses are minimized jointly. The primary loss applies to the patient-level logits against the patient-level label. The auxiliary loss applies to the visit-level logits against per-visit labels, with padding positions excluded. The total is the patient loss plus 0.3 times the visit loss. The auxiliary weight was chosen so that the visit-level signal guides learning without dominating; it prevents the model from ignoring intermediate visits and learning only from the final report, which would reduce HTT to Baseline behavior despite its access to the full sequence.

Optimization uses AdamW [13] at learning rate 2 *×* 10^*™*4^ with weight decay 0.01, applied only to trainable parameters: the LoRA matrices and all Level 2 components. A linear warmup spans the first 10% of steps, ramping from 10% of the target rate to the full value. A single NVIDIA RTX 5060 with 8 GB of VRAM set the batch arrangement: batch size 8 with gradient accumulation over 4 steps for an effective batch of 32, gradient-norm clipping at 1.0, and fp16 automatic mixed precision with a gradient scaler. Training ran for at most 20 epochs with early stopping on validation AUROC (patience 5), restoring the best-validation checkpoint for evaluation. Validation uses only training types in both experiments, so no held-out type influences any training decision. Table 5 lists every hyperparameter.

### 3.6 Evaluation protocol

AUROC is the primary metric because it does not move with class ratio. F1, precision, recall and, where relevant, specificity and accuracy are reported at the default threshold, taking the argmax of the SoftMax output. Following Mandrekar [14], AUROC between 0.80 and 0.90 is read as excellent and above 0.90 as outstanding.

In Experiment 1, HTT and the Baseline were trained on 280 patients, evaluated on 60 validation patients at each epoch, tested on 60 held-out patients at the end, and then evaluated on the 100 pancreatic patients never seen in training. In Experiment 2, the reference for transfer is the in-distribution test set of 660 reports, scored by the same checkpoint under the same conditions. Metrics are computed independently within each held-out type, and the reported mean is unweighted across the three.

## 4 Results

### 4.1 Experiment 1: temporal modelling on synthetic sequences

Table 6 and Figure 2a compare HTT and the Baseline on the synthetic corpus. Two evaluation sets are reported and they tell different stories, which we set out plainly.

**Table 6.**
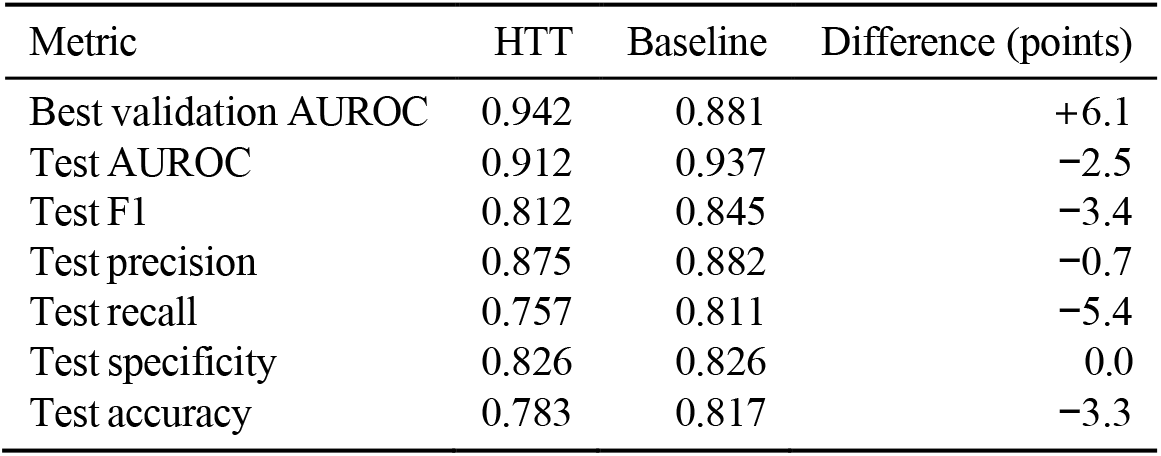
Experiment 1. In-distribution performance on synthetic sequential data. Validation AUROC is the checkpoint-selection criterion and is measured consistently across all epochs; the test set holds 60 patients, at which size a single reclassified patient shifts AUROC by roughly 1.5 to 2 points.

**Figure 2.**
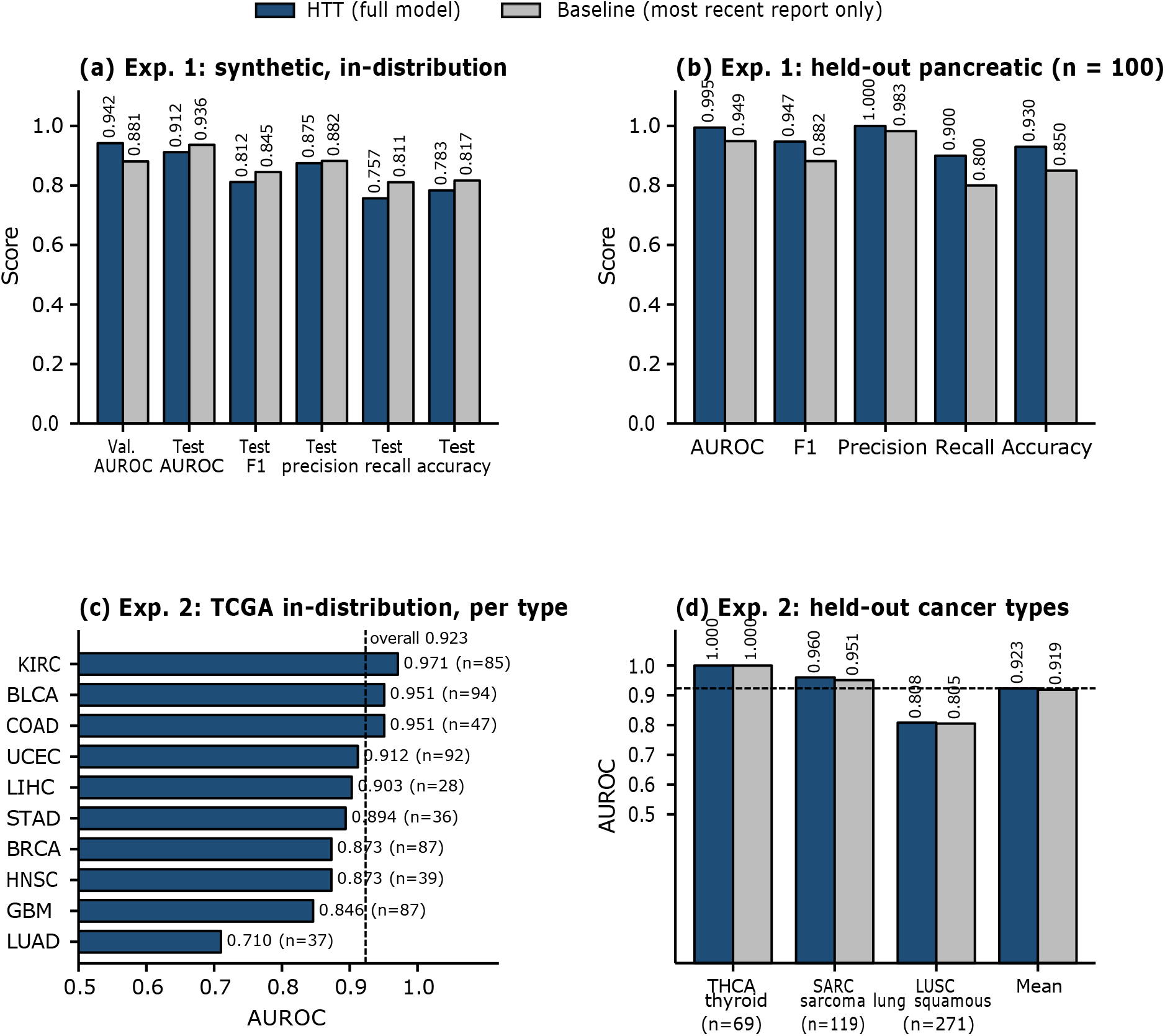
Summary of results. (a) Experiment 1, in-distribution: HTT against the Baseline on validation AUROC and on test-set metrics for 60 synthetic patients. (b) Experiment 1, held-out pancreatic cancer, 100 patients never seen in training. (c) Experiment 2, per-type AUROC on the TCGA in-distribution test set; the dashed line marks the overall test AUROC of 0.923 and PRAD is omitted because its AUROC is undefined. (d) Experiment 2, held-out AUROC for the three withheld cancer types and their unweighted mean, HTT against the Baseline; the dashed line marks the in-distribution test AUROC of 0.923.

On validation AUROC, measured at every epoch on the same 60 patients, HTT reaches 0.942 against 0.881 for the Baseline, a margin of 6.1 points. On the 60-patient test set the ordering reverses, with the Baseline at 0.937 and HTT at 0.912. At this sample size one reclassified patient moves AUROC by roughly two points, so the test comparison does not resolve a difference in either direction. What the experiment supports is a weaker statement than the validation margin alone would suggest: the temporal model reaches higher validation performance during training, and on held-in test data the two models are statistically indistinguishable.

Per-type performance on the synthetic test set (Table 7, 15 patients per type) ranges from AUROC 1.000 for breast cancer and 0.977 for NSCLC to 0.875 for CRC and 0.520 for prostate cancer. The near-chance prostate result most likely reflects the generator rather than the model: the prostate templates produced less discriminative trajectories than the others. Real prostate reports, with PSA values, Gleason scores and bone-scan findings, would present a different task.

**Table 7.** Experiment 1. HTT per-cancer-type performance on the synthetic test set, 15 patients per type.

| Cancer type | AUROC | F1 | $n$ |
| --- | --- | --- | --- |
| Breast cancer | 1.000 | 1.000 | 15 |
| Lung cancer (NSCLC) | 0.977 | 0.842 | 15 |
| Colorectal cancer | 0.875 | 0.833 | 15 |
| Prostate cancer | 0.520 | 0.600 | 15 |

#### Training dynamics

HTT’s validation AUROC oscillated between 0.42 and 0.66 for the first six epochs, then jumped from 0.659 to 0.898 at epoch 7 and reached 0.942 by epoch 10. This late, sharp improvement is consistent with hierarchical learning: the report encoder must first produce stable representations before the temporal transformer can find patterns across them.

#### Transfer to held-out pancreatic cancer

One hundred pancreatic patients were withheld entirely. Table 8 and Figure 2b report the result. HTT reaches AUROC 0.995 with no pancreatic training examples, against 0.949 for the Baseline, a margin of 4.6 AUROC points and 6.6 F1 points. The gap is larger and more consistent across metrics than on the in-distribution test set.

**Table 8.** Experiment 1. Transfer to held-out pancreatic cancer, *n* = 100 patients.

| Metric | HTT | Baseline | Difference (points) |
| --- | --- | --- | --- |
| AUROC | 0.995 | 0.949 | +4.6 |
| F1 | 0.947 | 0.882 | +6.6 |
| Precision | 1.000 | 0.983 | +1.8 |
| Recall | 0.900 | 0.800 | +10.0 |
| Accuracy | 0.930 | 0.850 | +8.0 |

The difference between the two models is concrete. The Baseline reads only the final report; if that report mentions new nodal involvement, it must decide from that phrase alone. HTT reads the whole sequence and knows that the first report described a stable 1.2 cm nodule, the second slight enlargement to 1.8 cm, and the third new nodal involvement. The continuous-time encoding adds the calendar distance between those observations, so a doubling over three months and a doubling over three years can be weighted differently. In this corpus the trajectory is encoded by construction, so the experiment shows that the architecture can read it; whether real clinical text carries trajectory signal of the same strength is a separate question the synthetic setting cannot answer.

### 4.2 Experiment 2: in-distribution performance on TCGA

Table 9 gives the overall result. On the 660-report test set spanning 11 cancer types, HTT reaches AUROC 0.923, F1 0.905, precision 0.881, recall 0.929 and accuracy 0.853.

**Table 9.** Experiment 2. HTT overall in-distribution performance on the TCGA test set, 11 cancer types, 660 reports.

| Metric | Value |
| --- | --- |
| Test AUROC | 0.923 |
| Test F1 | 0.905 |
| Test precision | 0.881 |
| Test recall | 0.929 |
| Test accuracy | 0.853 |
| Test specificity | 0.624 |

A single set of weights, with no per-type head and no per-type fine-tuning, ranks a random high-grade report above a random low-grade one about 92% of the time across a biologically diverse set of tumors.

Specificity is the weak point at 0.624. The corpus runs roughly 2.5:1 high to low grade and the model has absorbed that prior: it catches 93% of high-grade cases while correctly identifying only 62% of low-grade ones. For triage that asymmetry is arguably the right one, since missing an aggressive tumor cost more than over-flagging an indolent one, but the threshold would need recalibration for any use where false positives carry real cost. This is a further reason to treat AUROC, which is threshold-independent, as the primary summary.

Per-type performance (Table 10, Figure 2c) spans a wide band. Kidney renal clear cell carcinoma leads at AUROC 0.971, followed by bladder and colon at 0.951 each; uterine reaches 0.912 and liver 0.903. Glioblastoma sits at 0.846 and lung adenocarcinoma at 0.710, the weakest of the eleven. Prostate is undefined because every prostate report in the test split was high-grade, leaving no negative class; it is excluded from all averages.

**Table 10.** Experiment 2. HTT per-cancer-type performance on the TCGA in-distribution test set. PRAD AUROC is undefined because all prostate test samples were high-grade; it is excluded from averages.

| Code | Cancer type | AUROC | F1 | $n$ |
| --- | --- | --- | --- | --- |
| KIRC | Kidney renal clear cell carcinoma | 0.971 | 0.941 | 85 |
| BLCA | Bladder urothelial carcinoma | 0.951 | 0.989 | 94 |
| COAD | Colon adenocarcinoma | 0.951 | 0.919 | 47 |
| UCEC | Uterine corpus endometrial carcinoma | 0.912 | 0.904 | 92 |
| LIHC | Liver hepatocellular carcinoma | 0.903 | 0.700 | 28 |
| STAD | Stomach adenocarcinoma | 0.894 | 0.935 | 36 |
| BRCA | Breast invasive carcinoma | 0.873 | 0.884 | 87 |
| HNSC | Head and neck squamous cell carcinoma | 0.873 | 0.816 | 39 |
| GBM | Glioblastoma multiforme | 0.846 | 0.804 | 87 |
| LUAD | Lung adenocarcinoma | 0.710 | 0.889 | 37 |
| PRAD | Prostate adenocarcinoma | undefined | 1.000 | 28 |

### 4.3 Experiment 2: transfer to unseen cancer types

Table 11 and Figure 2d present the central result. No report from thyroid carcinoma, sarcoma or lung squamous cell carcinoma was used in training, validation or hyperparameter selection. HTT reaches AUROC 1.000 for thyroid carcinoma, 0.960 for sarcoma and 0.808 for lung squamous cell carcinoma, a mean of 0.923 across 459 reports. The in-distribution test AUROC is likewise 0.923, so there is no measurable degradation on transfer at these sample sizes.

**Table 11.**
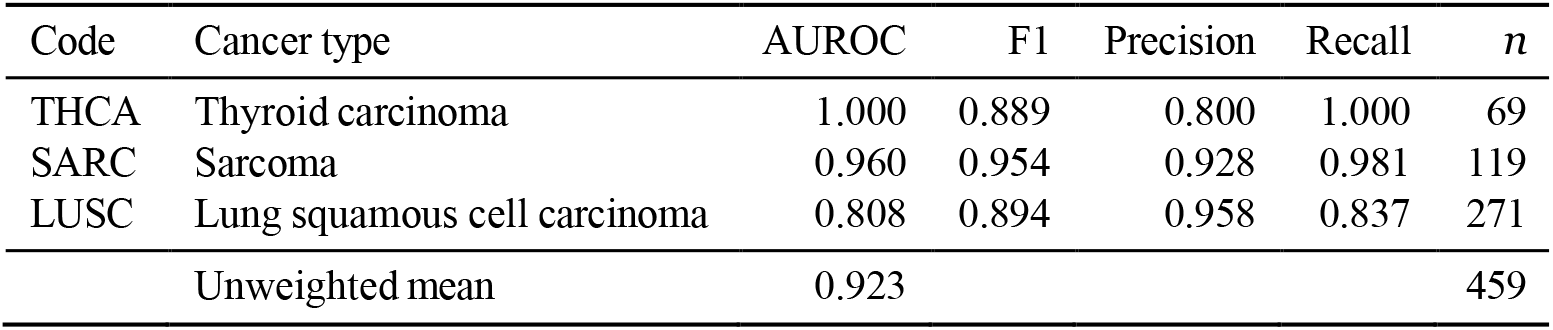
Experiment 2. Held-out performance on three cancer types absent from training, validation and model selection.

The size of this result is best read against the vocabulary gap. Thyroid grade is expressed through tall-cell variant histology, lymphovascular invasion and extrathyroidal extension. Sarcoma uses the FNCLCC score, whose three-component format appears nowhere in the training corpus. Lung squamous cell carcinoma is characterized by keratin pearl formation, intercellular bridges and nuclear-to-cytoplasmic ratios. A model dependent on the surface form of grade statements in the training types would transfer poorly to all three.

We attribute the transfer to the pre-trained encoder. BiomedBERT was pre-trained on a corpus in which descriptors such as “poorly differentiated”, “high mitotic rate” and “marked nuclear pleomorphism” occupy adjacent regions of representation space regardless of organ. Fine-tuning maps that shared region onto the grade objective, and the mapping holds for unseen types whose descriptors fall within the same region.

By the Mandrekar criteria [14], two held-out types are outstanding and the third excellent. The perfect thyroid AUROC is obtained on 69 reports and indicates complete separation within this set rather than a general guarantee; thyroid grading language is also comparatively explicit. Lung squamous cell carcinoma, at 0.808 on the largest held-out set, is the most informative case. It is the only held-out type whose anatomical site is represented in training, by lung adenocarcinoma, and shared vocabulary might have been expected to help. The opposite is observed. A plausible mechanism is representational interference: exposure to lung adenocarcinoma induces a lung-specific encoding of grade language partly misaligned with squamous criteria, which rest on keratinization and intercellular bridging rather than glandular architecture. Thyroid and sarcoma, lacking a competing prior, are evaluated through the general aggressiveness representation. Lung adenocarcinoma is also the weakest in-distribution type at 0.710. Establishing this mechanism would require holding out types at systematically varied pathological distances, which was not attempted here.

### 4.4 Ablation: what carries the transfer

Both models were trained on TCGA under identical conditions, so the ablation separates the encoder from the components layered above it. Table 12 reports the comparison. Mean held-out AUROC is 0.919 for the Baseline against 0.923 for HTT, a difference of 0.39 points, with per-type differences of 0.00, 0.89 and 0.30. All three are within the uncertainty of the sample sizes in Table 11.

**Table 12.**
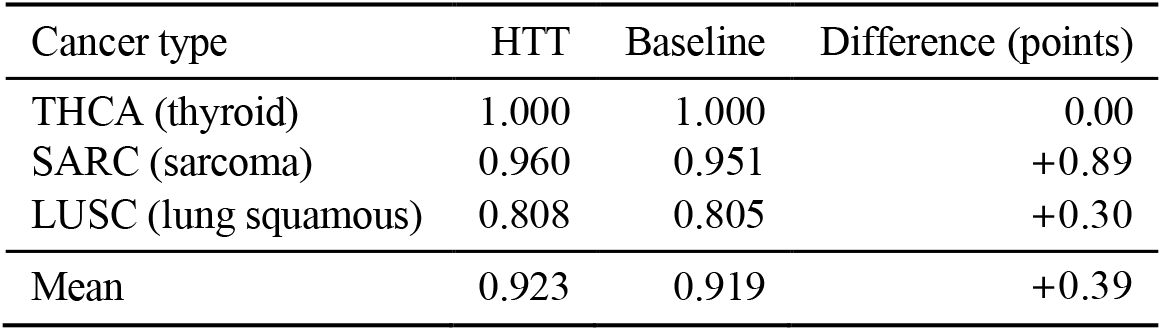
Experiment 2. Held-out AUROC, HTT against the non-temporal Baseline. Differences fall within the uncertainty of these sample sizes.

| Cancer type | HTT | Baseline | Difference (points) |
| --- | --- | --- | --- |
| THCA (thyroid) | 1.000 | 1.000 | 0.00 |
| SARC (sarcoma) | 0.960 | 0.951 | +0.89 |
| LUSC (lung squamous) | 0.808 | 0.805 | +0.30 |
| Mean | 0.923 | 0.919 | +0.39 |

This is the expected outcome and it is important to state why. TCGA-Reports holds one report per patient, so the temporal transformer operates on sequences of length one and reduces to a learned transform of a single conditioned embedding. Held-out types receive the reserved unseen-type index in place of a trained embedding, so type conditioning supplies little signal precisely where transfer is measured. Both facts bound what Level 2 can contribute in this experiment, and the ablation shows the bound is binding. The cross-cancer transfer observed on real data is therefore carried by the pre-trained encoder and the grade objective, not by the temporal components. Those components are exercised in Experiment 1, on the corpus that has sequences to read.

## 5 Discussion

### Why the held-out result is non-trivial

An AUROC of 1.000 on thyroid carcinoma, a type with its own grading vocabulary, anatomical context and histological features, shows the model has not memorized type-specific keywords. It has learned a representation of what pathological aggressiveness looks like in free text, and that representation extends to cancer families it never saw. The three held-out types were chosen precisely because their grading language differs from everything in training, and all three transferred.

### Implications for clinical deployment

Prevailing practice develops a separate model per cancer type. This scales poorly across a comprehensive cancer center and is infeasible for rare types. The transfer observed here indicates that a single model fine-tuned on a moderate number of common types can be applied to types absent from training, provided the target concept is already represented in the pre-trained encoder. Grade satisfies that condition. Extension to staging, margin status or treatment response is not supported by the present experiments.

### Pathological proximity and transfer

The ordering of held-out results runs against intuition. If representational interference explains it, then training-type selection for a transfer-oriented system cannot proceed by maximizing anatomical coverage alone, since a partially overlapping type may hurt performance on its histological neighbors. A design holding out histological variants of trained sites alongside unrelated types would test this directly.

### What the two experiments establish together

Experiment 1 shows that the temporal architecture can read trajectory information from a report sequence and that this helps most on a held-out cancer type, where the single-report model has least to go on. Experiment 2 shows that grade representations transfer across cancer families on real text, and that on a single-report corpus this transfer is carried by the encoder. Neither experiment alone demonstrates both capabilities on real longitudinal data, because no open corpus permits it. The design is the strongest available under that constraint, and the two results are consistent with one another rather than in tension.

### Clinical framing

An AUROC of 0.923 on unseen types supports assistive rather than autonomous use. The in-distribution specificity of 0.62 reflects the class prior of the training corpus, and threshold calibration against the relative costs of over- and under-calling would precede any deployment. Prioritization is the defensible application reports likely to describe aggressive disease are surfaced earlier for pathologist review, and the grade determination stays with the pathologist.

## 6 Limitations

### Synthetic data for temporal modelling

Experiment 1 rests on generated radiology text in which the progression label is inserted from the assigned trajectory. The corpus therefore cannot capture the linguistic complexity, abbreviation patterns or measurement specificity of real reports, and it cannot tell us whether real longitudinal text carries trajectory signal of comparable strength. Its results show that the architecture functions as designed on data whose ground truth is known exactly. They are a proof of mechanism, not an estimate of real-world performance, and the abstract numbers from this experiment should be read in that light.

### Validation against test in Experiment 1

The headline validation margin of 6.1 points is not reproduced on the 60-patient test set, where the Baseline is numerically ahead. We report both. The test set is too small to resolve the difference, and the validation set was the model-selection criterion, so neither number is a clean estimate. The held-out pancreatic result, on 100 patients the model never saw during training or selection, is the more reliable evidence from this experiment.

### Labels derived from pattern matching

TCGA ground truth was obtained by rule-based matching on report text. The approach is standard and was used in the original TCGA-Reports benchmark [11], but it admits label noise, and more consequentially the labels derive from the same text the model reads. A portion of the reported performance may reflect recovery of matched phrases rather than generalization. Two controls would bound this: a redacted condition replacing every matched span and its close variants with a neutral placeholder, and a bag-of-words logistic regression on the same text. Expert pathologist review of a stratified subsample would quantify residual label noise. All three are left to future work.

### Single report per patient in TCGA

The temporal component operates on sequences of length one in Experiment 2, so the two capabilities are demonstrated on separate corpora. A dataset with multiple longitudinal notes per patient, such as MIMIC-IV or GENIE BPC, would allow both to be tested on real data at once.

### Sample sizes

The held-out TCGA types comprise 69, 119 and 271 reports; the synthetic test set has 60 patients and the pancreatic set 100. The perfect thyroid AUROC and the sub-point ablation differences both carry wide uncertainty. Held-out types were chosen for diversity, not sampled systematically, so the mean should not be read as expected performance on an arbitrary unseen type.

### Encoder scale

All experiments use BiomedBERT-base under an 8 GB VRAM budget. Larger encoders may yield stronger representations, particularly for rare types, and evaluation of BioGPT-Large, Meditron-70B or BioMistral-7B on higher-memory hardware is left to future work.

## 7 Conclusion

This study presents the Hierarchical Temporal Transformer, a two-level architecture addressing two gaps in cancer NLP: the absence of temporal modelling and the absence of cross-cancer-type transfer. On a controlled synthetic corpus, the temporal model reaches a validation AUROC of 0.942 against 0.881 for a single-report baseline and transfers to held-out pancreatic cancer at 0.995 against 0.949, showing that the architecture can integrate trajectory information and real elapsed time in a setting where the ground truth is known. On 4,786 real TCGA pathology reports, the model predicts tumor grade for three cancer types withheld entirely from training at AUROC 1.000, 0.960 and 0.808, a mean of 0.923 that equals in-distribution performance. Ablation attributes this transfer to the pre-trained encoder and the grade objective.

The generalizable representation of pathological aggressiveness therefore appears to be formed during biomedical pre-training and recruited by fine-tuning. Progress on cross-type generalization for single-report corpora depends primarily on encoder capability and label fidelity, both constrained here and both addressable. The held-out-type protocol applies directly to any multi-type clinical corpus and gives a stricter generalization estimate than in-distribution testing. Biomedical language appears to generalize across tumor families, which supports unified pathology systems that require no per-type retraining.

## Data availability statement

The TCGA-Reports corpus [11] is openly available on the Hugging Face Hub. The synthetic data generator, label-extraction patterns, training configuration and evaluation scripts are available at https://github.com/rajneeshanand

The authors thank the TCGA-Reports curators for releasing the corpus in machine-readable form.

## Author contributions

N.B. and R.A. designed the model. R.A. implemented the training pipeline and ran the experiments. N.B. designed the evaluation protocol, performed the clinical interpretation and wrote the manuscript. H.H. contributed to the experimental design and revised the manuscript. D.Ç.S. supervised the work and revised the manuscript. All authors approved the final version.

## Conflicts of interest

The authors declare no conflicts of interest.

## A Glossary

Short definitions for readers on either side of the disciplinary boundary.

### AUROC

The probability that a randomly chosen positive case receives a higher model score than a randomly chosen negative case; 0.5 is chance, 1.0 is perfect ranking. Unaffected by class imbalance.

### F1 score

Harmonic mean of precision and recall.

### Pre-training and fine-tuning

Pre-training trains a model on a large general corpus; fine-tuning continues training on a task-specific dataset. BiomedBERT arrives fluent in medical language and needs only adjustment.

### Tokenization

Conversion of text into integer token IDs, with padding and truncation; each report is capped at 256 tokens.

### Embedding and mean pooling

The encoder maps each token to a 768-dimensional vector; mean pooling averages them into one vector per report.

### CLS token

A learnable token prepended to the sequence that aggregates it through attention; its output is the sequence representation.

### LoRA

Low-rank adaptation [8] inserts small trainable matrices beside frozen weights, so only 11.1% of parameters need gradients.

### Continuous-time positional encoding

Sinusoidal encoding applied to measured elapsed days rather than integer positions, so the model sees real calendar distances between visits.

### Unseen-type transfer

Evaluation on a cancer type with no examples in training or model selection.

### Tumor grade

A pathologist’s assessment of how abnormal tumor cells appear. High-grade (grade III, poorly differentiated) tumors grow and spread faster; low-grade (grade I, well differentiated) tumors resemble normal tissue.

### Cancer-type token

A marker such as [CANCER:BLCA] prepended to each report, giving an explicit cue and an index into the type embeddings.

### Ablation study

Removing a component to measure its contribution. Here the Baseline removes the temporal transformer, time encoding and type conditioning.

### Class imbalance

When one label dominates, accuracy misleads; AUROC does not.

## Notes

https://github.com/rajneeshanand

### Competing Interest Statement

The authors have declared no competing interest.

